# Identification of a putative RBOHD-FERONIA-CRK10-PIP2;6 plasma membrane complex that interacts with phyB to regulate ROS production in *Arabidopsis thaliana*

**DOI:** 10.1101/2025.11.23.689998

**Authors:** Devasantosh Mohanty, Yosef Fichman, María Ángeles Peláez-Vico, Ronald J Myers, Maya Sealander, Ranjita Sinha, Johanna Morrow, Ron Eckstein, Kate Olson, Chunhui Xu, Hong An, Chan Yul Yoo, Jian-Kang Zhu, Chunzhao Zhao, Sara I. Zandalinas, Emmanuel Liscum, Ron Mittler

**Affiliations:** Division of Plant Sciences and Technology, College of Agriculture Food and Natural Resources, Christopher S. Bond Life Sciences Center, University of Missouri; Columbia, MO 65211, USA; School of Plant Sciences and Food Security, Tel Aviv University, Ramat Aviv, Israel 6997801; Department of Biology and Environmental Sciences, Westminster College, Fulton, Missouri 65251; School of Biological Sciences, The University of Utah, Salt Lake City, UT 84112; Bioinformatics Analysis Core, Christopher S. Bond Life Sciences Center, University of Missouri; Columbia, MO 65211, USA; Institute of Advanced Biotechnology, Institute of Homeostatic Medicine, and School of Medicine, Southern University of Science and Technology, Shenzhen 518055, China; State Key Laboratory of Plant Trait Design, Shanghai Center for Plant Stress Biology, CAS center for Excellence in Molecular Plant Sciences, Chinese Academy of Sciences, Shanghai 200032, China; Department of Biology, Biochemistry and Environmental Sciences, University Jaume I, Av. de Vicent Sos Baynat, s/n, Castelló de la Plana 12071, Spain; Division of Biological Sciences, College of Arts & Sciences, University of Missouri, Columbia, MO, 65211, USA

**Keywords:** Feronia, Light stress, Phytochrome B, Reactive oxygen species (ROS), Respiratory burst oxygen homolog (RBOH)

## Abstract

Reactive oxygen species (ROS) regulate plant growth, development, and responses to the environment. ROS production by the RESPIRATORY BURST OXIDASE PROTEIN D (RBOHD) protein was recently shown to be regulated by PHYTOCHROME B (phyB), and phyB was found to be phosphorylated by FERONIA, highlighting the possibility that these three proteins interact to regulate ROS levels during stress.
Immunoprecipitation and proximity labelling, followed by split-luciferase and functional validation assays, were used to study the interactions between FERONIA, phyB, and RBOHD during excess light (EL) stress in *Arabidopsis thaliana*.
We reveal that phyB and FERONIA interact with RBOHD, that phosphorylation of phyB by FERONIA, as well as the kinase activity of FERONIA, are required for RBOHD-driven ROS production in response to EL stress, and that CYSTEINE-RICH RECEPTOR LIKE KINASE 10 (CRK10) and PLASMA MEMBRANE INTRINSIC PROTEIN 2;6 (PIP2;6) interact with RBOHD and phyB and are also required for EL-driven RBOHD ROS production.
Our findings uncover the existence of a putative plasma membrane complex between FERONIA, RBOHD, CRK10, and PIP2;6 that interacts with phyB to regulate ROS production in Arabidopsis in response to stress. This complex could play a canonical role in the integration and regulation of multiple signaling pathways in plants.

**Plain Language Summary:** We identified a complex between several different proteins at the plasma membrane that interacts with the light and temperature receptor protein phytochrome B to regulate reactive oxygen species formation during stress in plants. This complex could be involved in the regulation and integration of multiple abiotic and biotic signals in plants.

**ten:** ten

## INTRODUCTION

Reactive oxygen species (ROS), such as superoxide (O_2_^.-^) and hydrogen peroxide (H_2_O_2_), play a key role in the regulation of multiple stress, growth, and developmental processes in plants (Waszczak *et al*., 2018; Smirnoff & Arnaud, 2019; Considine & Foyer, 2021; Foyer & Hanke, 2022; Mittler *et al*., 2022; Peláez-Vico *et al*., 2024). Their steady-state levels in cells are determined by an interplay between different ROS production, scavenging, and transport mechanisms, that are transcriptionally, translationally and/or post-translationally regulated by various pathways, hormones, calcium, nitric oxide (NO) and hydrogen sulfide (H_2_S) (Smirnoff & Arnaud, 2019; Mittler *et al*., 2022). ROS can accumulate in different subcellular compartments in response to different biotic and/or abiotic stresses and their steady-state levels alter the redox state of cells, thereby control multiple signaling and metabolic pathways (Mittler & Jones, 2024). As part of their biological function in cells, ROS can directly alter the structure and thereby function of different proteins, channels, complexes, and enzymes, for example, by oxidizing cysteine or methionine residues, or by disrupting the function of iron-cluster centers (by O_2_^.-^ and H_2_O_2_, respectively; Young *et al*., 2019; Imlay *et al*., 2019; Mittler *et al*., 2022; Mittler & Jones, 2024; Karpinska & Foyer, 2024).

RESPIRATORY BURST OXIDASE HOMOLOGS (RBOHs) are among the most studied ROS producers of plant cells. They are plasma membrane (PM)-integral proteins that use NADPH at the cytosol to generate O_2_^.-^ at the apoplast. The generated O_2_^.-^ then dismutates spontaneously or enzymatically (catalyzed by superoxide dismutases; SODs) to H_2_O_2_ at the apoplast and diffuses back into the cytosol via aquaporins (Suzuki *et al*., 2011; Luo *et al*., 2021; Yu *et al*., 2023, 2024; Arnaud *et al*., 2023; Goto *et al*., 2024a; Zhan *et al*., 2025; Chen *et al*., 2025). In *Arabidopsis thaliana*, the RBOH proteins D and F (RBOHD and RBOHF, respectively) regulate multiple developmental, growth, reproduction, and stress responses at the different aboveground parts of plants, while the RBOH protein C (RBOHC), as well as other root localized RBOHs, regulate ROS production and root responses belowground. Among the well-studied roles of RBOHs in plants are the regulation of stomatal responses to changes in environmental conditions (*e.g.,* Postiglione & Muday, 2020), the regulation of plant defense mechanisms in response to pathogen infection (*i.e.,* the oxidative burst; Kadota *et al*., 2015), and the regulation of systemic signaling in response to different abiotic or biotic stimuli (*i.e.,* the ROS wave; Fichman & Mittler, 2020a).

At the base of RBOH regulation in cells are multiple transcriptional, post-translational, and protein-protein mechanisms. RBOHs were found to physically interact with different kinases, such as BOTRYTIS-INDUCED KINASE 1 (BIK1) that together with different phosphatases phosphorylate/de-phosphorylate RBOHs and regulate their enzymatic activities (Goto *et al*., 2024b). In addition, RBOHs interact with other proteins such as Rho-of-Plants (ROPs), and calcium/different calcium binding proteins (Smokvarska *et al*., 2020). These interactions, as well as nitrosylation, persulfidation, acetylation, recycling via clathrin mediated endocytosis, and/or other biochemical modifications, alter RBOHs function and allow the turning of ROS production from ‘high’ to ‘low’ and *vice versa* in response to different stimuli (Smith *et al*., 2014; Lee *et al*., 2022; Zhang *et al*., 2025). As RBOHs function can be attenuated or amplified via multiple modifications and/or interactions (triggered by different converging pathways; *e.g.,* pathways involved in biotic and/or abiotic stresses, development, and growth), RBOHs are thought to integrate multiple pathways and fine-tune the overall process of ROS accumulation in cells (Mittler *et al*., 2022).

Two other multi-signal integrators and master regulators of cells, that have been recently linked to ROS production in plants, are the PM-localized FERONIA receptor kinase (Duan *et al*., 2022; Liu *et al*., 2023, 2024b,a; Xu *et al*., 2024; Jiang *et al*., 2024; Tang & Guo, 2025), and the cytosolic-nuclear light receptor and transcriptional regulator PHYTOCHROME B (phyB; Han *et al*., 2019, 2024; Devireddy *et al*., 2020; Pierik & Ballaré, 2021; Hernando *et al*., 2021; Quail, 2021; Fichman *et al*., 2023; Ma *et al*., 2023; Viczián & Nagy, 2024)). While FERONIA was found to be involved in ROS production during pathogen and abiotic stress responses, phyB was shown to be required for ROS production in response to excess light stress, as well as several other abiotic stress conditions (but not wounding). The regulation of ROS production by phyB during abiotic stresses was further linked to RBOHD, suggesting that these two proteins could interact (Fichman *et al*., 2023), and FERONIA was recently shown to phosphorylate phyB and regulate responses of Arabidopsis to salt stress (Liu *et al*., 2023). Taken together, the studies described above suggest that phyB, RBOHD, and FERONIA use a common axis of ROS signaling regulation and could even interact with each other.

To test the hypothesis that FERONIA, phyB and RBOHD interact to regulate ROS production in plants, we conducted proximity labeling and protein complex immunoprecipitation for phyB and RBOHD, respectively, followed by *in vivo* split-luciferase and functional ROS production (in response to excess light stress) assays, as well as *in vitro* and *in silico* validation experiments. We found that phyB and FERONIA interact with RBOHD, that phosphorylation of phyB by FERONIA, as well as the kinase function of FERONIA, are required for RBOHD-driven ROS production in response to excess light (EL) stress, and that CYSTEINE-RICH RECEPTOR LIKE KINASE 10 (CRK10) and PLASMA MEMBRANE INTRINSIC PROTEIN 2;6 (PIP2;6) interact with RBOHD and phyB and are also required for EL-driven RBOHD ROS production. Our findings reveal therefore the existence of a putative PM complex between FERONIA, RBOHD, CRK10, and PIP2;6 that interacts with phyB to regulate ROS production in Arabidopsis in response to EL stress. As several key signal integrators are involved in the complex identified here (*i.e.,* RBOHD, FERONIA, and phyB), and each of these is absolutely required for ROS production in cells in response to EL stress, the complex we identified could play a canonical role in the integration and regulation of multiple signaling pathways in plants.

## RESULTS

### Identification and validation of protein interactors of RBOHD during excess light stress

To identify putative proteins that interact with RBOHD during EL stress, we tagged RBOHD with an HA-tag at its N-terminal and expressed it under the control of the native *RbohD* promoter in the *rbohd* mutant (Fig. **1a**). In agreement with our previous studies showing that RBOHD is responsible for ROS production in Arabidopsis in response to EL stress (Fichman *et al*., 2019, 2023; Zandalinas *et al*., 2020), the expression of the HA-tagged RBOHD in the *rbohd* mutant could restore ROS production in response to a 10 min whole plant EL stress exposure (Fig. **1b**). To identify proteins that interact with RBOHD in response to EL stress, we subjected *rbohd* and transgenic *rbohd* plants expressing the HA-tagged RBOHD (*rbohd*/*RbohD*:HA-RBOHD; Fig. **1c**) to control (no stress), an EL stress of 10 min (EL), and dark conditions (Dark), and processed them for immunoprecipitation using antibodies against the HA tag bound to Sepharose beads. As RBOHD is an PM integral protein, we obtained membrane-enriched fractions of control, EL-treated, and dark-adapted *rbohd* and *rbohd*/HA-RBOHD plants before conducting immunoprecipitation. As shown in Fig. **1d** and Table S1, our HA-RBOHD immunoprecipitation analysis identified FERONIA (*At3g51550*), CRK10 (*At4g23180*), and PIP2;6 (*At2g39010*) as interacting with RBOHD under all 3 conditions (Control, EL stress, and Dark). While CRKs were found to regulate ROS production and stress responses in plants (Kimura *et al*., 2020; Piovesana *et al*., 2023), PIPs are thought to mediate the transport of H_2_O_2_ across membranes in both plants and animals (Medraño-Fernandez *et al*., 2016; Prado *et al*., 2019; Mukherjee *et al*., 2024; Jia *et al*., 2025). In addition to CRK10, FERONIA and PIP2;6, our analysis identified several other interesting proteins such as CALMODULIN 7 (*At3g43810*) and CALCIUM-BINDING PROTEIN CML13 (*At1g12310*), WALL-ASSOCIATED RECEPTOR KINASEs 2 (*At1g21270*) and 8 (*At1g16260*), several LEUCINE-RICH REPEAT (LRR) like receptors (*At1g53430*, *At3g14840*, *At1g33590*, *At3g20820*, *At5g63410*, and *At5g07910*), SERINE/THREONINE-PROTEIN KINASE BSK3/4/7 (*At4g35230*, *At1g01740*, and *At1g63500*), SERINE/THREONINE-PROTEIN PHOSPHATASE 2A (*At3g25800*), BRASSINOSTEROID INSENSITIVE 1 (*At4g39400*), and several proteins involved in RBOHD recycling via clathrin mediated endocytosis (*At3g11130*, *At5g46630*, *At3g08530*, *At2g25430*, and *At2g40060*). Several of these proteins (or their homologs) were previously reported to interact with RBOHD supporting the validity of our study (Fichman & Mittler, 2020a; Lee *et al*., 2022; Zhang *et al*., 2025). The interactions of the proteins highlighted above with RBOHD should be addressed in future studies as they may shed additional light on the different abiotic/EL stress/biotic-associated interactors of RBOHD. Below, we focused on FERONIA, CRK10 and PIP2;6.

**Figure 1.**
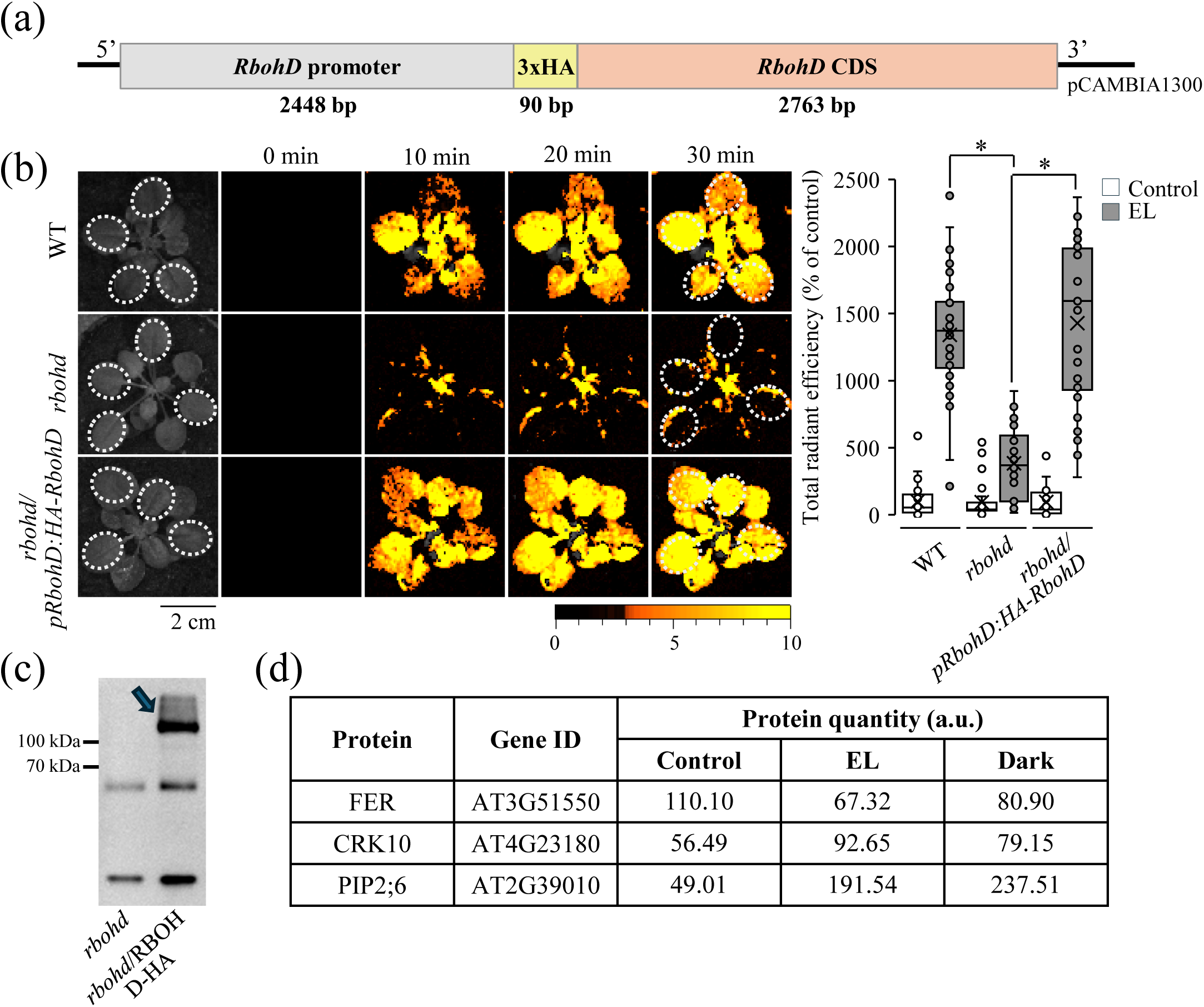
Identification of proteins that interact with RBOHD during excess light stress. (**a**) A scheme of the HA-RBOHD construct used to transform the *rbohd* mutant and conduct immunoprecipitation. (**b**) Representative time-lapse whole-plant reactive oxygen species (ROS) images (Left) and a bar graph showing quantification of the ROS signal (Right) of wild type (WT), *rbohd*, and *rbohd* plants complemented with the HA-tagged RBOHD protein driven by the endogenous *RbohD* promoter (a), in response to a whole-plant 10 min excess light (EL) stress. (**c**) A protein blot showing that the HA-tagged RBOHD protein driven by the endogenous *RbohD* promoter is expressed in transgenic plants. (**d**) A table showing the detection of FERONIA (FER), CRK10, and PIP2;6 in the HA-tag based immunoprecipitation of RBOHD. Confirmation of protein-protein interactions are shown in Figs. 2 and 3. A complete list of all RBOHD putative interactors is shown in Table S1. Statistical significance was determined by a Student’s *t*-test: \**P* < 0.05 (N≥3). Abbreviations: a.u., arbitrary unit; bp, base pairs; cm, centimeter; CDS, coding DNA sequence; CRK10, Cysteine-Rich Receptor-like Kinase 10; EL, excess light; FER, FERONIA; min, minutes; HA, hemagglutinin; kDa, kilo Dalton; PIP2;6, Plasma membrane Intrinsic Protein 2;6; RbohD, Respiratory burst oxygen homologue D; WT, Wild type.

To confirm that FERONIA, CRK10 and PIP2;6 interact with RBOHD *in vivo*, we conducted split-luciferase assays between the cytosolic kinase domains of FERONIA or CRK10, or the entire PIP2;6 protein, and the N-or C-terminals of RBOHD, that are cytosolic facing (Zhang *et al*., 2025). As shown in Fig. **2**, the cytosolic kinase domains of FERONIA or CRK10, or the entire PIP2;6 protein, interacted with the N-terminal domain of RBOHD. In contrast, no interactions were observed for these domains, or the entire PIP2;6 protein, with the C-terminal of RBOHD.

**Figure 2.**
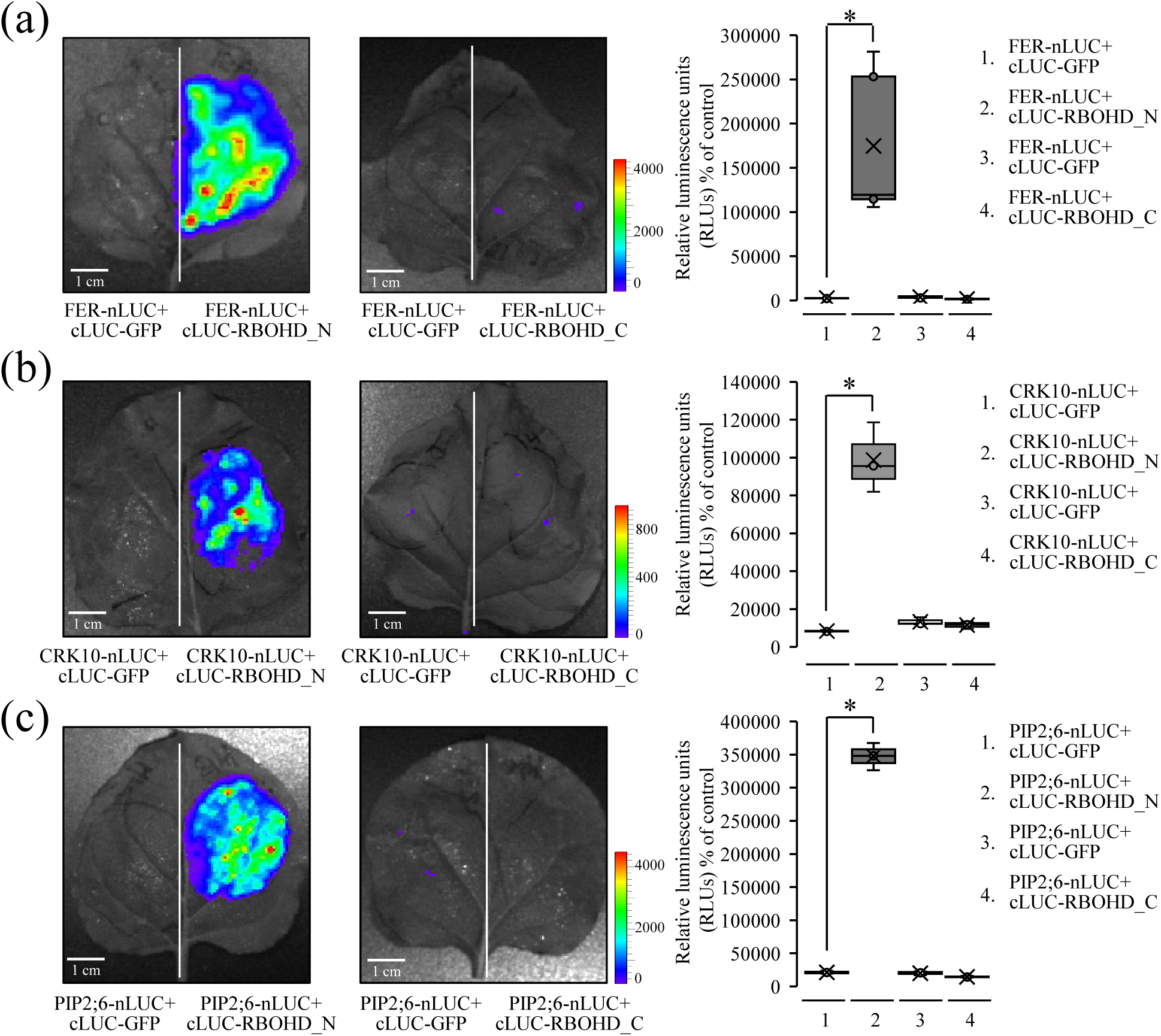
Split-luciferase *in vivo* protein-protein interaction studies between RBOHD (N-and C-terminals) and FERONIA, CRK10, and PIP2;6. (**a**) Representative images (Left), and a bar graph showing quantification of the luciferase signal (Right), indicating interactions between RBOHD N-terminal and FERONIA (FER). (**b**) Same as in (a), but for the interactions between RBOHD N-terminal and CRK10. (**c**) Same as in (a), but for the interactions between RBOHD N-terminal and PIP2;6. Statistical significance was determined by using Student’s *t*-test: \**P* < 0.05 (N≥3). Abbreviations: cm, centimeter; cLuc, C-terminal Luciferase; CRK10, Cysteine-Rich Receptor-like Kinase 10; FER, Feronia; GFP, Green Fluorescent Protein; nLuc, N-terminal Luciferase; PIP2;6, Plasma membrane Intrinsic Protein 2;6; RLU, Relative luminescence unit; RbohD_N, Respiratory burst oxygen homologue D N-terminal; RbohD_C, Respiratory burst oxygen homologue D C-terminal.

To determine whether the interactions between RBOHD and CRK10, PIP2;6, or FERONIA are essential for ROS production in response to EL stress, we conducted whole-plant ROS imaging (Fichman *et al*., 2019) analyses of wild type (WT) plants and different mutants deficient in the function of these proteins. As shown in Fig. **3**, in contrast to WT plants, two independent mutants for CRK10 (*crk10*; Fig. **3a**) or PIP2;6 (*pip2;6*; Fig. **3b**) did not accumulate ROS in response to a 10 min whole plant EL stress exposure. As shown in Fig. **3c**, similar results were found for a mutant of FERONIA (*fer4*). To further test whether the kinase domain of FERONIA is essential for ROS production in response to EL stress, we used the *fer4* mutant complemented with the FERONIA protein expressed under its native promoter (*fer4/FER*), as well as the *fer4* mutant complemented with a dead kinase (DK) FERONIA protein expressed under its native promoter (*fer4/FER-DK*; Liu *et al*., 2023). As shown in Fig. **3c**, expression of the WT FERONIA protein (*fer4/FER*), but not the dead kinase FERONIA protein (*fer4/FER-DK*) in the *fer4* mutant was able to complement ROS production in response to a 10 min whole plant EL stress.

**Figure 3.**
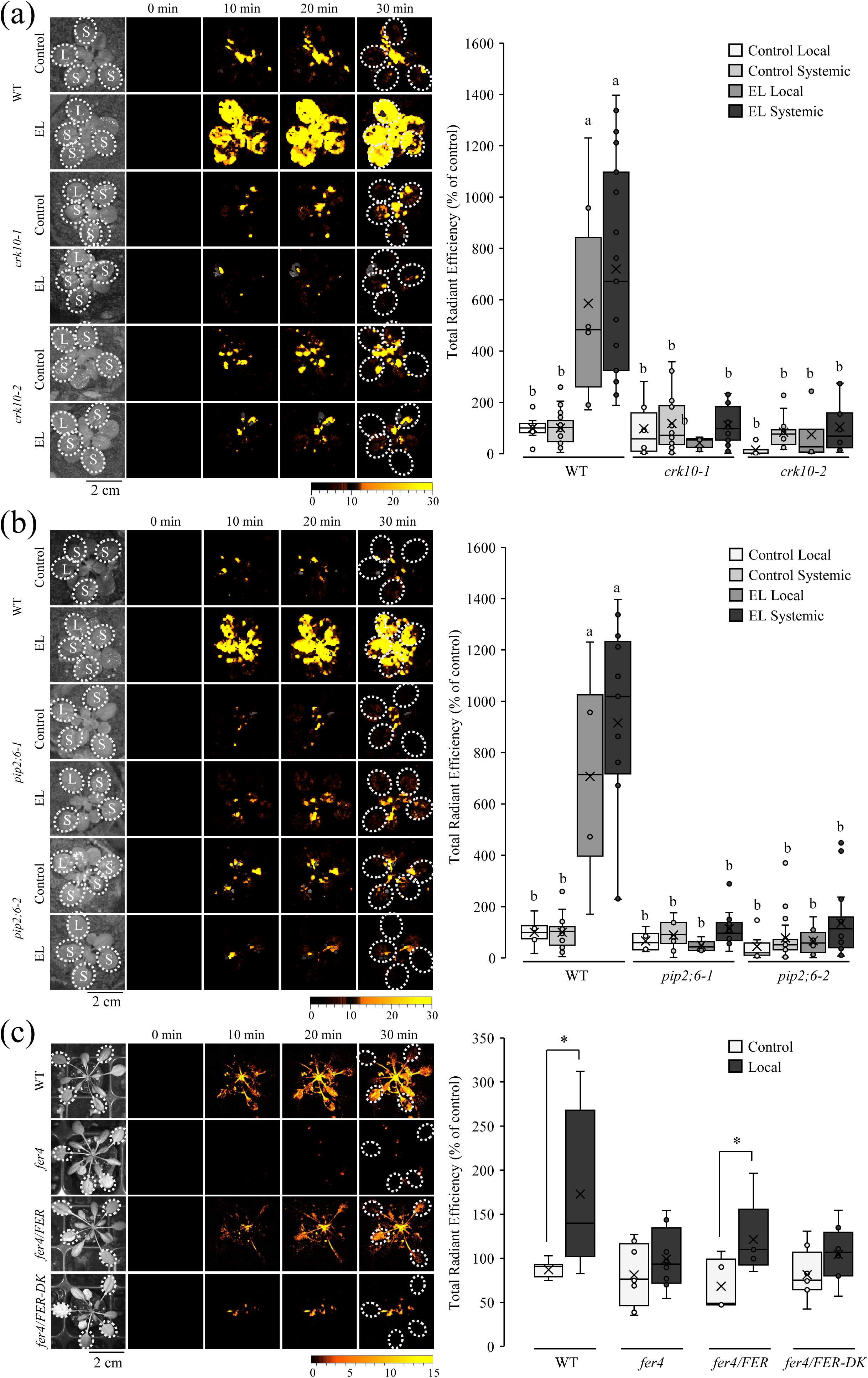
Whole-plant reactive oxygen species (ROS) imaging analyses of CRK10, PIP2;6 and FERONIA mutants subjected to excess light stress. (**a**) Representative time-lapse images (Left), and a bar graph showing quantification (Right), of whole-plant ROS levels in wild type (WT) and two independent mutants for CRK10 (*crk10-1*, *crk10-2*), subjected to a whole-plant 10 min EL stress. (**b**) Same as in (a), but for two independent mutants of PIP2;6 (*pip2;6-1*, *pip2;6-2*). (**c**) Representative time-lapse whole-plant ROS images (Left), and a bar graph showing quantification of the ROS signal (Right), of wild type (WT), a mutant of FERONIA (*fer4*), a *fer4* mutant complemented with the WT FERONIA protein (*fer4/FER*), or a *fer1* mutant complemented with a ‘dead kinase’ (DK) version of the FERONIA protein (*fer4/FER-DK*). Statistical significance was determined by using Student’s *t*-test: \**P* < 0.05 (N≥3), or one-way ANOVA (Fisher Least Significant Difference; Different letters above bars indicate significance at P<0.05; N=3). Abbreviations: cm, centimeter; CRK10, Cysteine-Rich Receptor-like Kinase 10; DK, dead kinase; EL, excess light; FER, Feronia; L, local leaf; min, minutes; PIP2;6, Plasma membrane Intrinsic Protein 2;6; S, systemic leaf; WT, wild type.

Taken together, our findings support a model in which FERONIA, CRK10 and PIP2;6 interact with RBOHD and are required for EL-induced ROS production by this key regulatory protein (Figs. **1-3**). In addition, we show that the kinase activity of FERONIA is essential for ROS production in response to EL stress in Arabidopsis (Fig. **3c**).

### Identification and validation of protein interactors of phyB during excess light stress

To identify proteins that interact with phyB during EL stress, we used TurboID-mediated proximity labelling (Feng *et al*., 2024). For this purpose, we used transgenic *phyAphyB* plants expressing the phyB coding sequence fused to a miniTurboID protein under the control of the ubiquitin promoter (*phyAphyB/UBQ10p:PhyB-miniTurbo*; Fig. **4a**; Olson *et al.,* 2025). In agreement with our previous studies showing that phyB is essential for ROS production in response to EL stresses in Arabidopsis (Fichman *et al*., 2023), the expression of the phyB fused to the miniTurboID protein was able to restore ROS production in the *phyAphyB* or *phyB* backgrounds in response to a 10 min whole plant EL stress exposure (Figs. **4b** and **S1**). To identify proteins that interact with phyB in response to EL stress, we treated *phyAphyB/UBQ10pPhyB-miniTurbo* plants with biotin, subjected them to control or 10 min EL stress, and processed them for biotin-based proximity labelling. As our goal was to identify proteins that are in proximity to phyB, as well as RBOHD, and RBOHD is a PM integral protein, we obtained membrane-enriched fractions of biotin-treated control and EL stress-treated *phyAphyB/UBQ10p:PhyB-miniTurbo* plants, before conducting proximity labelling enrichment and detection via proteomics analysis (Fig. **4c**). As shown in Fig. **4d** and Table S2, our phyB-miniTurbo proximity analysis of the membrane-enriched fractions identified RBOHD, FERONIA, CRK10, and PIP2;6 as interacting with phyB under control or EL stress conditions (Fig **4d**; Table S2). The identification of RBOHD, FERONIA, CRK10, and PIP2;6 as phyB interactors supported our HA-RBOHD immunoprecipitation results (Figs. **1-3**; Table S1), as well as our previous work proposing a functional link between phyB and RBOHD (Fichman *et al*., 2023), revealing that FERONIA, CRK10, and PIP2;6 could interact with both phyB and RBOHD. While no PHYTOCHROME-INTERACTING FACTORS (PIFs) were identified in our proximity labelling analysis (likely due to our focus on the membrane-enriched fractions), several other interesting proteins were identified as potentially interacting with phyB. These included CRYPTOCHROME 1 (*At4g08920*), LESION SIMULATING DISEASE 1 (LSD1) zinc finger family protein (*At4g20380*), PHYTOCHROME C and E (*At5g35840* and *At4g18130*), PHYTOCHROME KINASE SUBSTRATE 2 (*At1g14280*) and phytochrome-associated PROTEIN PHOSPHATASE TYPE 2C (*At1g22280*), multiple receptor proteins [*e.g.,* RECEPTOR LECTIN KINASE, RECEPTOR LIKE PROTEIN 29, PROLINE-RICH RECEPTOR-LIKE KINASE 2C, several LRR-like receptors, CALMODULIN-BINDING RECEPTOR-LIKE CYTOPLASMIC KINASE 2, and CRK17; *At2g37710*, *At2g42800*, *At3g01560*, *At1g33590*, *At1g33600*, *At1g49750*, *At1g58848*, *At1g62630*, *At3g15410*, *At3g20820*, *At3g24480*, *At4g13340*, *At4g18670*, *At4g19510*, *At4g19530*, *At5g12940*, *At5g43470*, *At5g45510*, *At5g48620*, *At5g61240*, *At5g66900*, *At4g00330*, and *At4g23250*, and BRI1-ASSOCIATED RECEPTOR KINASE (*At4g33430*)]. These putative interactors should be addressed in future studies as they may shed more light on the different membrane-associated interactors of phyB. An interaction between phyB and PM-localized proteins was previously reported (Jaedicke et al., 2012; Liu *et al*., 2023), further supporting our findings that phyB could interact with the PM-localized protein RBOHD.

**Figure 4.**
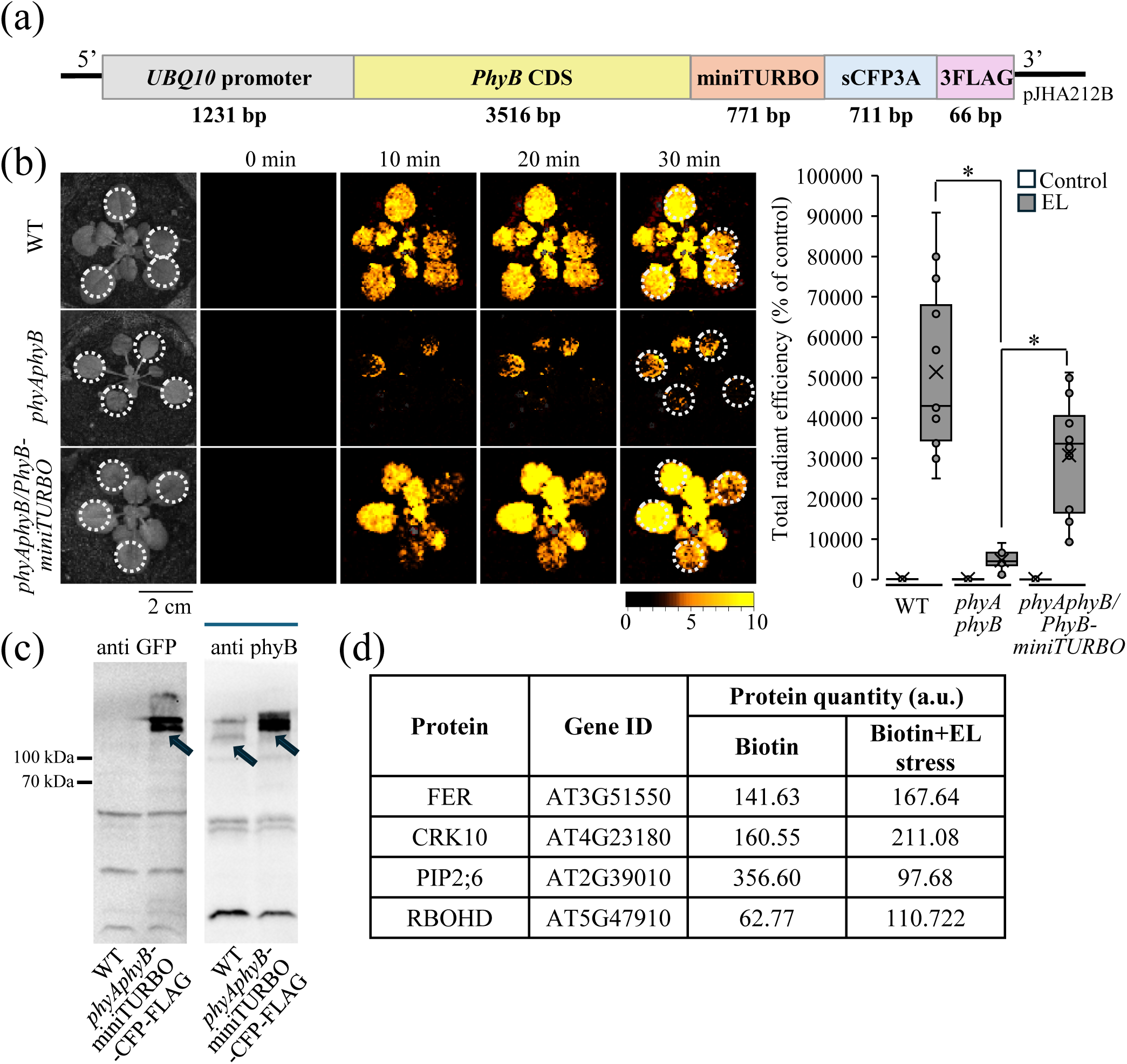
Identification of proteins that interact with PHYTOCHROME B (phyB) during excess light stress. (**a**) A scheme of the phyB construct used to transform the *phyAphyB* mutant and conduct proximity labeling. (**b**) Representative time-lapse whole-plant reactive oxygen species (ROS) images (Left) and a bar graph showing quantification of the ROS signal (Right) of wild type (WT), *phyAphyB* and *phyAphyB* plants complemented with the phyB-miniTurbo protein driven by the ubiquitin 10 (*UBQ10*) promoter (a), in response to a whole-plant 10 min excess light (EL) stress. (**c**) Protein blots showing that the phyB-miniTurbo protein driven by the *UBQ10* promoter is expressed in transgenic plants. (**d**) A table showing the detection of FERONIA (FER), CRK10, PIP2;6, and RBOHD in the proximity labelling analysis of phyB. Confirmation of protein-protein interactions are shown in Figs. 5 and 7. Complementation of the *phyB* mutant by the phyB-miniTurbo protein is shown in Fig. **S1**. Statistical significance was determined by using Student’s *t*-test: * *P* < 0.05 (N≥3). A complete list of all putative phyB interactors is shown in Table S1. Abbreviations: a.u., arbitrary unit; bp, base pairs; cm, centimeter; CDS, coding DNA sequence; CFP, Cyan Fluorescent Protein; CRK10, Cysteine-Rich Receptor-like Kinase 10; EL, excess light; FER, Feronia; kDa, kilo Dalton; min, minutes; phyAphyB, PhytoschromeAPhytochromeB; PIP2;6, Plasma membrane Intrinsic Protein 2;6; RbohD, Respiratory burst oxygen homologue D; WT, Wild type.

To validate the interactions of RBOHD, FERONIA, CRK10 and PIP2;6 with phyB *in vivo*, we conducted split-luciferase assays between phyB (entire protein) and the cytosolic kinase domains of FERONIA or CRK10, the cytosolic N-and C-terminals of RBOHD, or the entire PIP2;6 protein. As shown in Fig. **5a**, phyB interacted with both the N-and C-terminals of RBOHD. phyB also interacted with the cytosolic domains of CRK10 and FERONIA (**Figs. 5b** and **5c**, respectively), and with PIP2;6 (Fig. **5d**). In addition, as shown in Fig. **S2**, we performed *in vitro* immunoprecipitation analysis of phyB and RBOHD N-or C-terminals. This analysis revealed that phyB can interact with the N-or C-terminals of RBOHD *in vitro*, further supporting our proximity and split-luciferase analyses (Figs. **4** and **5**).

**Figure 5.**
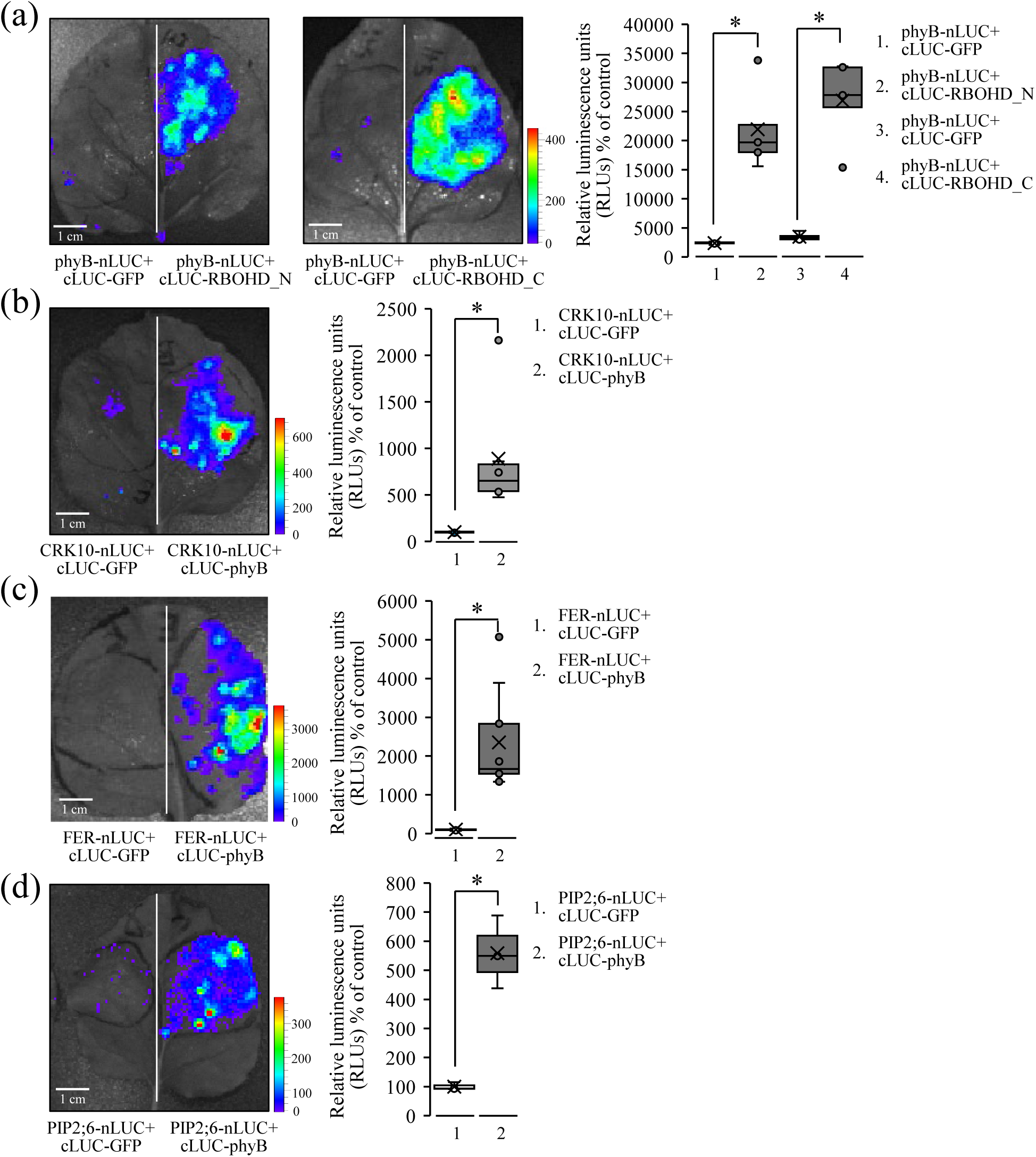
Split-luciferase *in vivo* protein-protein interaction studies between PHYTOCHROME B (phyB) and RBOHD (N- and C- terminals), FERONIA, CRK10, and PIP2;6. (**a**) Representative images (Left), and a bar graph showing quantification (Right), of the luciferase signal indicating interactions between phyB and the RBOHD N- or C- terminals. (**b**) Same as in (a), but for the interactions between phyB and CRK10. (**c**) Same as in (a), but for the interactions between phyB and FERONIA. (**d**) Same as in (a), but for the interactions between phyB and PIP2;6. Statistical significance for all bar graphs was determined by using Student’s *t*-test: \**P* < 0.5 (N≥3). Abbreviations: cm, centimeter; cLuc, C-terminal Luciferase; CRK10, Cysteine-Rich Receptor-like Kinase 10; FER, Feronia; GFP, Green Fluorescent Protein; nLuc, N-terminal Luciferase; PIP2;6, Plasma membrane Intrinsic Protein 2;6; phyB, Phytochrome B; RLU, Relative luminescence unit; RbohD_N, Respiratory burst oxygen homologue D N-terminal; RbohD_C, Respiratory burst oxygen homologue D C-terminal.

### Phosphorylation of phyB by FERONIA is required for ROS production in response to EL stress

FERONIA was previously shown to interact with and phosphorylate phyB at positions S106 and S227 and to regulate salt tolerance in Arabidopsis (Liu *et al*., 2023). Our current analysis revealed that FERONIA interacts with phyB and RBOHD and that its kinase activity is required for ROS production in response to EL stress [Figs. **1-3**; In addition, we previously reported (Fichman *et al*., 2023) as well as currently show (Fig. **4b**), that phyB is also required for ROS production in response to EL stress]. Taken together, these findings suggest that FERONIA could interact with and phosphorylate phyB during EL stress, and that this phosphorylation is required for ROS production in response to this abiotic stress. To test this hypothesis, we subjected WT, *phyB* mutants, and *phyB* mutants complemented with the WT phyB protein (*phyB*::35S:phyB-GFP), a phyB protein that cannot be activated by FERONIA via phosphorylation (mutations S106A and S227A in phyB; *phyB*::35S:phyB^(S106A, S227A)^-GFP; Liu *et al*., 2023), or a *phyB* mutant complemented with a constitutively active phyB protein (mutations S106D and S227D in phyB; *phyB*::35S:phyB^(S106D, S227D)^-GFP; Liu *et al*., 2023), to a whole-plant EL stress of 10 min and imaged ROS production. As shown in Figs. **6** and **S3**, expression of the WT phyB protein in the *phyB* mutant restored ROS production in response to EL stress, while expression of the constitutively active phyB protein (S106D and S227D; *phyB*::35S:phyB^(S106D, S227D)^-GFP) in the *phyB* mutant caused high levels of ROS to be produced in response to EL stress. In contrast to the complementation of the *phyB* mutant with the WT or constitutively active phyB proteins, expression of the phosphorylation inactive phyB protein (S106A and S227A; *phyB*::35S:phyB^(S106A, S227A)^-GFP) in the *phyB* mutant could not complement the EL stress-induced ROS production phenotype. Taken together, these findings support our hypothesis that FERONIA binds to, phosphorylates, and activates phyB, and that this phosphorylation is required for ROS production by RBOHD in response to EL stress (Fig. **6**; Liu *et al*., 2023).

**Figure 6.**
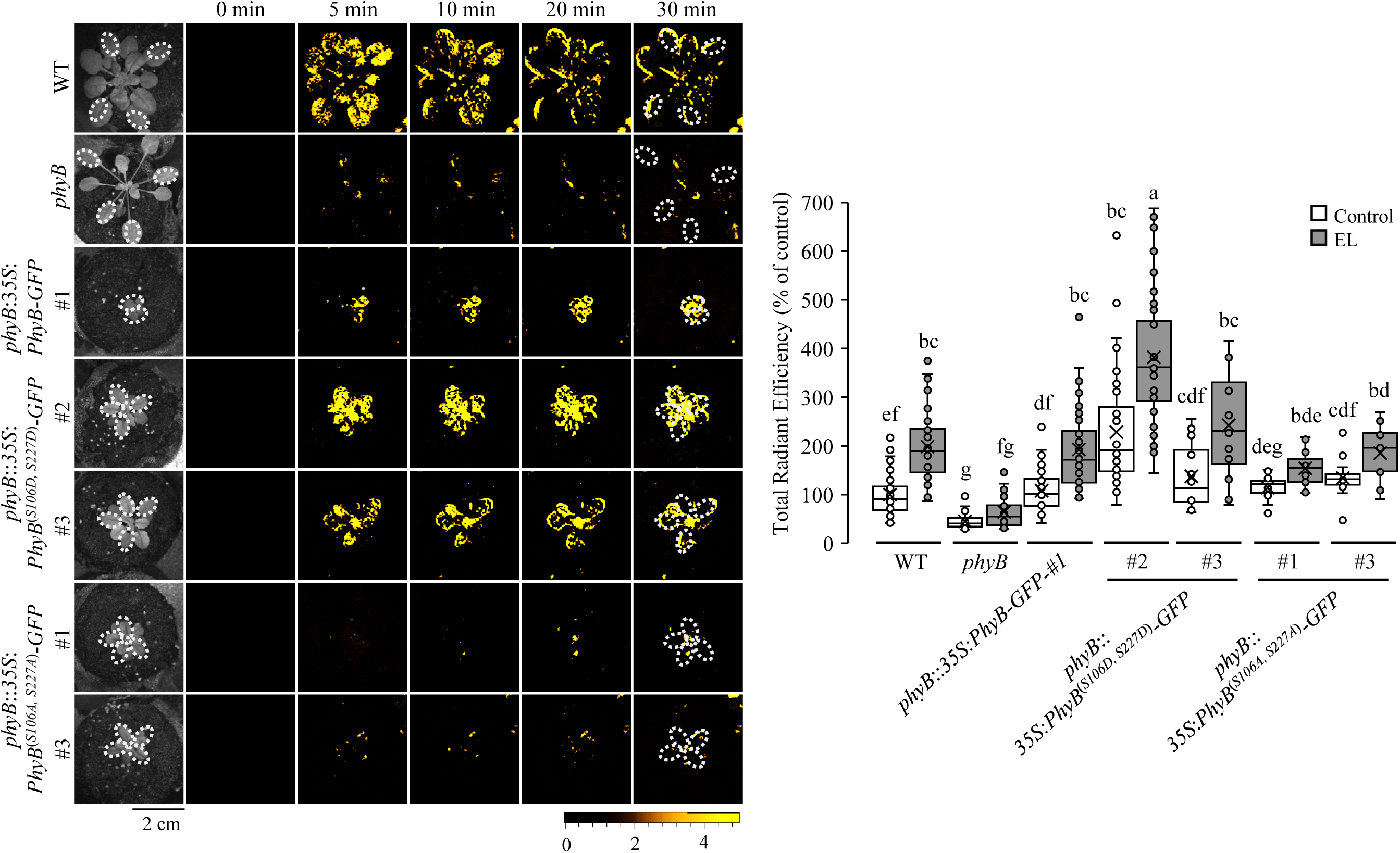
Whole-plant reactive oxygen species (ROS) imaging analyses of the PHYTOCHROME B mutant (*phyB*) complemented with wild type phyB or mutants of phyB. Representative time-lapse whole-plant ROS images (Left), and a bar graph showing quantification (Right) of the ROS signal for wild type (WT), the *phyB* mutant, the *phyB* mutant complemented with the phyB protein expressed under the control of the *CaMV35* promoter (*phyB*:3*5S*:*PhyB-GFP*), the *phyB* mutant complemented with a phyB protein that mimics constitutive phosphorylation of phyB [*phyB*:*PhyB^(S106D/S227D)^-GFP*], or the *phyB* mutant complemented with a phyB protein that cannot be phosphorylated by FERONIA [*phyB*:*PhyB^(S106A/S227A)^-GFP*]. Statistical significance was determined by one-way ANOVA (Fisher Least Significant Difference; Different letters above bars indicate significance at P<0.05; N≥3). Abbreviations: A, Alanine; cm, centimeter; D, Aspartic acid; EL, excess light; GFP, Green Fluorescence Protein phyB, Phytochrome B; cm, centimeter; S, Serine; WT, Wild type.

### *In silico* modeling of phyB-RBOHD-CRK10-FERONIA-PIP2;6 interactions

To further determine whether a complex between phyB, RBOHD, CRK10, FERONIA, and PIP2;6 is structurally feasible, we conducted *in-silico* modeling of the putative interactions between these different proteins. Such analysis provides a statistical confidence score for the different interactions between each pair of proteins, as well as an opportunity to visualize them and ascertain that they occur in a logical manner (*e.g.,* with respect to the PM and orientation of the different proteins imbedded in it). We first examined the interactions between phyB and the N- or C-terminals of RBOHD, the cytosolic domains of FERONIA and CRK10, or the entire PIP2;6 protein. As shown in Fig. **7a**, phyB interacted with the cytosolic domains of RBOHD, CRK10 and FERONIA with a confidence score of 0.95-0.88 (Table S3), suggesting that these interactions are feasible. In line with its cytosolic-nuclear localization, phyB interacted with parts of PIP2;6 that are outside the PM with a confidence score of 0.99 (Fig. **7a**; Table S3). When RBOHD interactions with FERONIA, phyB, CRK10, and PIP2;6 were modeled *in-silico*, confidence scores were also high (0.99-0.96; Fig. **7b**; Table S4). Interestingly, when the interactions between all 5 proteins was modeled (Fig. **8a**) it was found that the transmembrane domains of PIP2;6, FERONIA, CRK10 and RBOHD were all aligned, that the apoplastic domains of FERONIA and CRK10 interacted on one side of the transmembrane domains, and that phyB interacted with the cytosolic domains of RBOHD, CRK10 and FERONIA on the other side of the transmembrane domains. This arrangement (Fig. **8a**) corresponded with membrane orientation and protein localization of the 5 proteins involved in the complex (Fig. **8b**).

**Figure 7.**
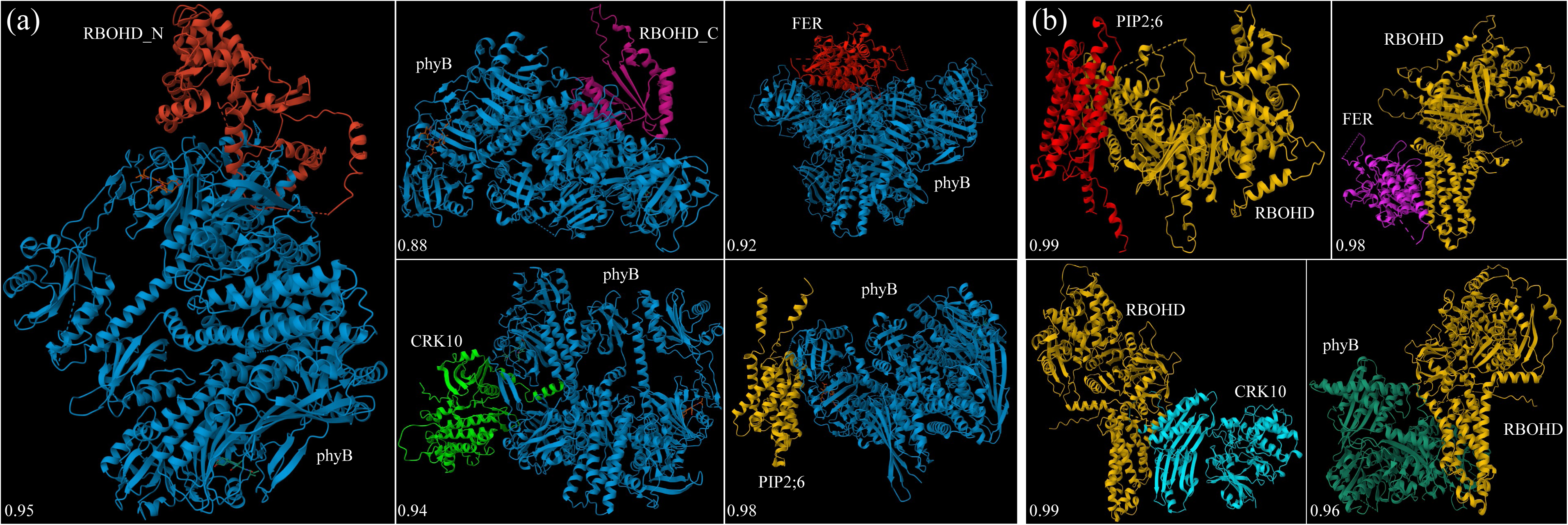
*In silico* modeling of protein-protein interactions between the different proteins proposed to form a complex at the plasma membrane. (**a**) Modeling of the putative interactions between the entire phyB protein and RBOHD N- or C- terminals, the cytosolic domains of FERONIA or cRK10, or the entire PIP2;6 protein. (**b**) Modeling of the putative interactions between RBOHD, PIP2;6, phyB, or the cytosolic domains of FERONIA or CRK10. Scores for best fit of each model are given on the bottom left corner of each model. Protein folding was predicted by AlphaFold and pairwise interaction models were generated using the HDOCK server. Abbreviations: CRK10, Cysteine-Rich Receptor-like Kinase 10; FER, Feronia; phyB, Phytochrome B; PIP2;6, Plasma membrane Intrinsic Protein 2;6; RbohD-C, Respiratory burst oxygen homologue D C-terminal; RbohD-N, Respiratory burst oxygen homologue D N-terminal.

**Figure 8.**
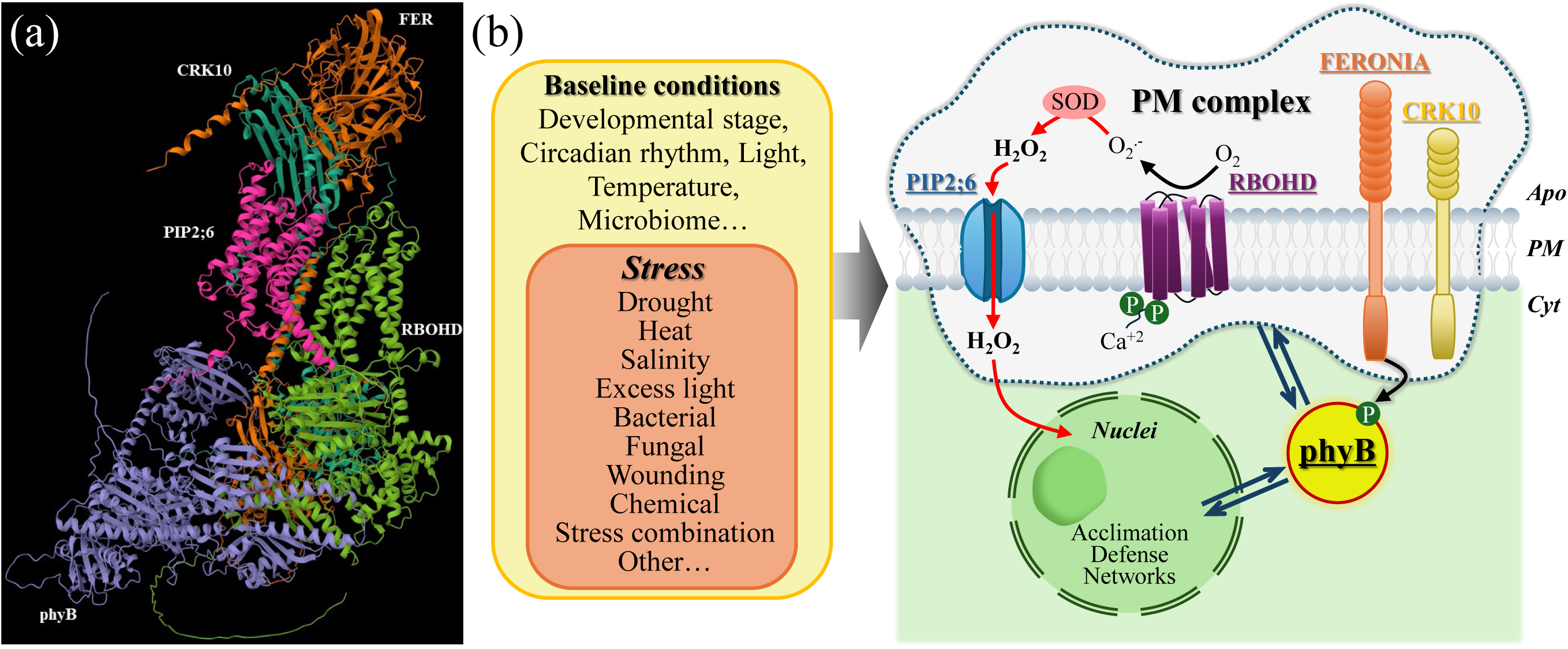
*In silico* analysis of the entire complex and a model for its putative function. (**a**) *In silico* modeling of the putative interactions between all members of the complex showing that all transmembrane domains are aligned, that the apoplastic portions of FERONIA and CRK10 interact on one side of the transmembrane domains, and that all cytosolic domains of the different proteins interact on the other side of the transmembrane domains. (**b**) A hypothetical model for the putative complex. FERONIA is proposed to phosphorylate phyB and this phosphorylation is shown to be important for reactive oxygen species (ROS) production by RBOHD. Hydrogen peroxide produced as a result of RBOHD activation is proposed to enter the cell through PIP2;6 and together with phyB alter nuclear transcription of different acclimation and defense pathways. CRK10 is also shown to be required for ROS production by RBOHD. Protein folding was predicted by AlphaFold and pairwise interaction models were generated using the HDOCK server. Abbreviations: Apo, apoplast; CRK10, Cysteine-Rich Receptor-like Kinase 10; Cyt, cytoplasam; FER, Feronia; phyB, Phytochrome B; PIP2;6, Plasma membrane Intrinsic Protein 2;6; PM, plasma membrane; RbohD, Respiratory burst oxygen homologue D; SOD, Superoxide Dismutase.

## DISCUSSION

We previously hypothesized that phyB interacts with RBOHD to regulate ROS production in response to EL stress in Arabidopsis (Fichman *et al*., 2023). Here, we used *in vivo*, *in vitro*, and *in silico* tools to demonstrate that such an interaction occurs in Arabidopsis and that it plays a key role in regulating ROS production in response to EL stress. Interestingly, we identified several other proteins that interact with phyB and RBOHD and could form a functional complex at the PM of plants (Fig. **8**). In particular, we identified FERONIA that is a PM localized receptor kinase, recently shown to phosphorylate phyB and regulate plant responses to salinity (Liu *et al*., 2023). Using a mutant of FERONIA with a dead kinase domain (Fig. **3c**), and mutants of phyB that cannot be phosphorylated by FERONIA (Fig. **6**), we further show that phyB phosphorylation by FERONIA could be essential for ROS production under conditions of EL stress. Taken together, these findings suggest a model in which RBOHD and FERONIA interact at the PM with phyB to regulate ROS production in response to stress (Fig. **8b**). In line with this model, all three proteins were required for ROS production during EL stress (Figs. **1**, **4**, **6**), and the phosphorylation of phyB by FERONIA was required for this process (Fig. **6**). In addition to RBOHD and FERONIA, we also identified CRK10 and PIP2;6 as involved in this putative complex and required for ROS production during EL stress (Figs. **1**, **3**, **4**). CRK2 was previously shown to phosphorylate RBOHD on its C-terminal during responses to pathogen infection (Kimura *et al*., 2020), and PIP2;1 was shown to be required for ROS production during EL stress (Fichman *et al*., 2021). It is therefore possible that CRK10 phosphorylates RBOHD in response to EL stress, and that PIP2;6 (or heteromeric complexes between PIP2;6 and other PIPs, some identified by our proximity labelling with phyB, Table S2; or with PIP2;1 identified in Fichman et al., 2021) mediate the transfer of H_2_O_2_ generated in the apoplast by apoplastic or cell wall bound SODs (*e.g.,* Vanacker *et al*. 1998; Kukavica *et al*. 2009; Wang *et al*., 2014; García *et al*., 2020; Chen *et al*., 2022) into the cytosol (Jia *et al*., 2025). The proximity of PIP2;6 to RBOHD could generate an efficient RBOHD generator of O_2_^.-^, that is converted to H_2_O_2_ in the apoplast by SODs, for cytosolic (or even nucleus) signaling purposes (Fig. **8b**). Further studies are needed to substantiate these roles of CRK10 and PIP2;6. An intriguing possibility that also arises from our study is that RBOHD could generate superoxide radicals, and that due to the acidic conditions at the apoplast and the proximity of RBOHD to PIP2;6, some of these superoxide radicals can become protonated and available for transport. Protonated superoxide could potentially transfer through PIPs or other channels at the PM (Fisher, 2009; Cordeiro, 2015; Karpinska & Foyer, 2024) into the cytosol, become deprotonated due to the more basic cytosolic pH and then dismutated into H_2_O_2_ by cytosolic CuZnSOD. In this respect it should be noted that our proximity labelling of phyB identified MnSOD1 (usually mitochondrial) and FeSOD1 (usually chloroplastic) as interacting with phyB. However, if this intriguing possibility is correct, it would most likely be mediated by cytosolic CuZnSODs. It should also be noted that all 5 proteins identified by our study are expressed in mature leaves (Table S5), further supporting our hypothesis that they interact to regulate ROS production during EL stress in this tissue (Figs. **1**, **3**, **4**).

While ROS could be produced via multiple pathways during EL stress (*e.g.,* chloroplasts, mitochondria, and peroxisomes (Waszczak *et al*., 2018; Smirnoff & Arnaud, 2019; Considine & Foyer, 2021; Foyer & Hanke, 2022; Mittler *et al*., 2022; Peláez-Vico *et al*., 2024), we previously provided evidence that ROS production during EL stress, applied using our EL method, and measured by our ROS imaging analysis, is primarily mediated by RBOHD and regulated by phyB (Fichman *et al*., 2019, 2023). These findings are supported here by our findings that EL stress induced ROS production that is suppressed in the *rbohd* mutant, could be complemented by expressing RBOHD in the *rbohd* mutant background (Fig. **1b**), and that ROS production that is suppressed in the *phyB* mutant, could be complemented by expressing phyB in the *phyB* mutant (Fig. **4b**). The putative complex we identified in this study most likely therefore regulates RBOHD-produced ROS during EL stress (Fig. **8b**).

Studies of phyB function in cells are mainly focused on its role as a light or temperature sensor that regulates transcriptional responses in the nuclei. Our previous (Devireddy *et al*., 2020; Fichman *et al*., 2023) and current work, as well as work of several other labs (*e.g.,* Jaedicke *et al*., 2012; Liu *et al*., 2023) has revealed however that phyB may have several additional roles in cells that are not linked to transcriptional regulation in the nuclei. As we previously shown, phyB is required for ROS production in response to EL stress, cold, heat, or pathogen infection (Fichman *et al*., 2023). phyB is also required for salinity responses in Arabidopsis (Liu *et al*., 2023). To mediate these roles, phyB could function as a stress/light receptor and/or a scaffold protein that is essential for multiple biotic and abiotic ROS-mediated responses in plants, inside or outside the nuclei. The identification of FERONIA and CRK10 as associated with phyB and RBOHD at the PM, highlights the cytosolic role(s) of phyB and links phyB through these receptors to multiple abiotic and/or biotic stimuli. Thus, the putative complex we identified in this study could constitute a major integrator of different abiotic and/or biotic signals/stimuli leading to enhanced or suppressed ROS production in cells. As ROS levels regulate multiple processes in plants (Waszczak *et al*., 2018; Smirnoff & Arnaud, 2019; Considine & Foyer, 2021; Foyer & Hanke, 2022; Mittler *et al*., 2022; Peláez-Vico *et al*., 2024), this complex could play a key role in the regulation and integration of growth, development, and responses to different environmental stimuli, and phyB could be playing a key role in these processes. In future studies it would be interesting to explore whether different stress conditions alter the context of the putative complex we identified and whether other important receptors and channels (*e.g.,* HPCA1 and/or MSL3; Fichman *et al*., 2022) are associated with it.

## MATERIAL AND METHODS

### Plant materials and growth conditions

*Arabidopsis thaliana* cv. Columbia-0 wild-type (WT), *rbohd* (*At5g47910*; Torres *et al*., 2002), *phyB-9* (*At2g18790*; Reed *et al.,* 1993), *phyA-211/phyB-9* (Reed *et al.,* 1994), *pip2;6-1* (*At3g39010*, SALK_118213C), *pip2;6-2* (SALK_092140C), *crk10-1* (*At4g23180*, SALK_116653C), *crk10-2* (SALK_017511C), *fer4* (*At3g51550*, Haruta *et al.,* 2018), *rbohd/RbohDp:3xHA-RbohD*, *phyAphyB/UBQ10p:phyB CDS – miniTurbo-sCFP3A-3FLAG* (Olson *et al.,* 2025), *phyB/35S:PhyB-GFP*, *phyB/35S:PhyB^(S106D, S227D)^-GFP*, and *phyB/35S:PhyB^(S106A, S227A)^-GFP* (Liu *et al*., 2023) plants were grown under a 10-h light (80 μmol photons m^-2^ s^-1^)/14-h dark regime at 20°C, 65% relative humidity. The light cycle started at 08:00 AM and ended at 06:00 PM. Plants were grown on peat pellets (Jiffy 7; Jiffy International, Kristiansand, Norway) for 4 to 5 weeks. T-DNA insertion mutants used in this study were obtained from the Arabidopsis Biological Resource Center (ABRC; https://abrc.osu.edu/). All T-DNA mutants were genotyped by allele-specific PCR to identify homozygous mutant plants for the respective insertion. Primers used to genotype these mutants are listed in Table S6.

### Application of fluorescent probes, stress treatment, and image acquisition

For ROS imaging, four-to five-week-old *Arabidopsis* plants were fumigated in a glass chamber (24.5 x 12.5 x 16.5)” with 10 μM DCFH-DA (2′,7′-Dichlorofluorescin diacetate; excitation/emission-480 nm/520 nm; Catalog number: D6883, SIGMA-ALDRICH, Saint Louis, MO, USA) prepared in 10 mL of 10 mM phosphate buffer (pH7.4) containing 0.002% Silwet L-77 (LEHLE seeds, Round Rock, TX, USA) using a portable mini nebulizer (Punasi Direct, Hong Kong, China) for 30 minutes as described (Fichman *et al*., 2019, 2022; Fichman & Mittler, 2020b, 2021). Excess light (EL) stress was applied by exposing whole plants to 740 μmol photons m^−2^ s^−1^ light for 10 minutes using a LED array light source (Bestva, Commerce, CA, USA) as described (Fichman *et al*., 2019). Plants were imaged using the IVIS Lumina S5 platform and Living Image 4.7.3 software in acquisition mode (PerkinElmer, Waltham, MA, USA) via a constant image set (excitation/emission 480 nm/520 nm). Images were captured every 60 s for 30 minutes as described (Fichman *et al*., 2019; Fichman & Mittler, 2020b, 2021; Myers *et al*., 2023), and total radiant efficiency of regions of interest (ROIs) were calculated as described (Mohanty *et al*., 2025). Each experiment was repeated at least three times with four to five technical repeats, and data from all technical repeats of each experiment were pooled to generate the final graph.

### *In vivo* co-immunoprecipitation and proteomics analysis

Five-week-old *rbohd* mutant and *rbohd/RbohDp:3xHA-RbohD* homozygous plants were used to perform *in vivo* co-Immunoprecipitation. Plants were kept under control conditions, subjected to complete darkness overnight, or exposed to EL stress (740 μmol photons m^−2^ s^−1^) for 10 minutes. One g leaf samples were harvested and immediately frozen in liquid nitrogen. Frozen leaf samples were ground in pre-chilled pestle and mortar using 5 mL of homogenization buffer [50 mM HEPES (pH7.5), 250 mM sucrose, 5% glycerol, 10 mM EDTA, 0.5% PVP, 50 mM NaPP, 1 mM NaMo, 25 mM NaF, 1X protease inhibitor cocktail (Catalog number: A32955, Pierce Protease Inhibitor Mini Tablets, EDTA-Free; Lee *et al*., 2020)]. Cell debris were removed by filtering with Miracloth followed by centrifugation at 8,000g for 10 minutes at 4°C. The supernatant was centrifuged using an ultracentrifuged at 100,000g for 30 minutes at 4°C to enrich for membranes. Pellets were washed twice with 1 mL of homogenization buffer. The membrane fraction was solubilized using homogenization buffer supplemented with 2% IGEPAL (Catalog number: CA-630, SIGMA-ALDRICH, Saint Louis, MO, USA) for 30 minutes at 4°C. Post solubilization the membrane fraction was diluted 10 times using 3 mL of IP Buffer (150 mM Tris-HCl pH7.5, 150 mM NaCl, 10% glycerol, 10 mM DTT, 1 mM PMSF, 1X completer protease inhibitor). 20 µL Protein A-Sepharose beads per sample (Catalog number: P9424, SIGMA-ALDRICH, Saint Louis, MO, USA) were equilibrated by washing 5 times with IP buffer at 4°C. The solubilized membrane fractions were precleared by incubating with Protein A-Sepharose beads for 90 minutes at 4°C on a space rotor. Post preclearing the membrane fractions were harvested using a centrifuged at 2,000 RPM for 2 minutes at 4°C and incubated on a space rotor overnight with 30 µL of equilibrated Protein A-Sepharose beads per sample and 2.5 µg of anti-HA antibody (Catalog number: 66006-2-Ig, Proteintech, Rosemont, IL, USA) at 4°C. Post incubation the beads were harvested using a centrifuged at 2,000 RPM for 2 minutes at 4°C and washed 3 times with IP Buffer supplemented with 0.2% IGEPAL. Complete removal of detergent was ensured by 3 further washes with IP buffer. Beads were resuspended in 30 µL of 1x SDS loading buffer [31.5 mM Tris-HCl (pH6.8), 10% glycerol, and 1% SDS] and incubated at 70°C for 10 minutes to elute the proteins. 2 µL of the elute were used for SDS-PAGE and western blot analysis and rest of the samples were submitted for LCMS analysis. IP proteins were precipitated from the SDS-PAGE buffer with acetone. All samples were digested with trypsin. Peptides (entire reaction) were loaded onto Evotips (C18 tips, Evosep) and processed according to the manufacturer’s protocol (Evosep). Peptides were analyzed by mass spectrometry as follows: peptides were separated using the Evosep One 60SPD method (22 minutes) on a Pepsep15 analytical column (Bruker 15 cm x 150 µm x 1.5 µm ReproSil-PurC18). The Evosep One HPLC was attached to a Bruker timsTOF-PRO mass spectrometer via a Bruker CaptiveSpray source. MS data were collected in positive-ion data-independent acquisition PASEF mode (Meier *et al.,* 2020) over an m/z range of 300.5 to 1349.5 and a mobility range of 0.65 to 1.45. Fifty DIA windows were collected (of varying m/z width) for a total cycle time of 2.76 sec. An ion-mobility-based collision energy was used with 20 eV at 0.85 and 59 eV at 1.30. The raw data was copied to Spectronaut V18.9 and searched against the Uniprot-Arabidopsis database using the directDIA utility employing the Pulsar algorithm. Proteins were identified (single-peptide protein hits were ignored) based on trypsin digestion (2 missed cleavages were allowed), carbamidomethyl-Cys as fixed modification, oxidized Met and AcetylK as variable mods, 20 ppm precursor mass tolerance, 0.1 Da fragment mass tolerance.

### Split luciferase complementation assay

Four-week-old *Nicotiana benthamiana* plants were used to perform *Agrobacterium*-mediated transient co-expression of proteins fused to N-or C-terminal of Luciferase to study protein-protein interactions. *Nicotiana* plants were grown on soil at 28°C under 14/10-h light/dark cycle. Full length PhyB, cytosolic domain of CRK10 (aa348-634), full length PIP2-6, cytosolic domain of FER (aa470-895) were cloned in pCAMBIA1300-nLuc between the *Kpn*I and *Sal*I sites. RBOHD N-terminus (aa1-376), RBOHD C-terminus (aa756-922), full length PhyB were cloned in pCAMBIA1300-cLuc between the *Kpn*I and *BamH*I sites. Primers used to generate these constructs are listed in Table S5. The constructs were electroporated into *Agrobacterium tumefaciens* strain GV3101 and transient co-expression studies were performed in *Nicotiana* plants to study protein-protein interactions as described (Zhou *et al*., 2018). *Agrobacterium* transformed with *35S*:*cLuc-GFP* was used as control in appropriate combination. 72 h post infection *Nicotiana benthamiana* plants were fumigated inside a glass chamber with 1 mM D-luciferin potassium salt (Catalog number: ab143655, Abcam, Waltham, MA, USA) prepared in 10 mL of 10 mM phosphate buffer (pH7.4) containing 0.002% Silwet L-77. Post fumigation the leaves were imaged using the IVIS Lumina S5 platform (PerkinElmer, Waltham, MA, USA). The relative luminescence units of regions of interest (ROIs) were calculated using the Living Image 4.7.3 software (PerkinElmer).

### TurboID-based *in vivo* proximity labeling and proteomics analysis

Five-week-old *phyAphyB/UBQ10p:PhyB CDS – miniTurbo-sCFP3A-3FLAG* homozygous transgenic plants were fumigated with 10 mL of 200 µM biotin (Catalog number: 58-85-5, SIGMA-ALDRICH, Saint Louis, MO, USA) prepared in 10 mM phosphate buffer (pH7.4) containing in 0.002% Silwet L-77 (or just buffer) using a portable mini nebulizer for 30 minutes in a glass chamber. Post fumigation plants were exposed to EL stress (740 μmol photons m^−2^ s^−1^) for 10 minutes. Post stress exposure samples were immediately collected and frozen in liquid nitrogen. One mg of frozen leaf samples were ground in pre-chilled pestle and mortar using 5 mL of homogenization buffer (50 mM HEPES (pH7.5), 250 mM sucrose, 5% glycerol, 10 mM EDTA, 0.5% PVP, 50 mM NaPP, 1 mM NaMo, 25 mM NaF, 1X protease inhibitor cocktail; Lee *et al*., 2020). Cell debris were removed by filtering with Miracloth followed by centrifugation at 8,000g for 10 minutes at 4°C. The supernatants were centrifuged using an ultracentrifuged at 100,000g for 30 minutes at 4°C to enrich for membranes. The pellets were washed twice with 1 mL of homogenization buffer to remove excess biotin. The membrane fraction was solubilized using homogenization buffer supplemented with 5% SDS for 30 minutes at room temperature. Post solubilization the membrane fraction was diluted 10 times using RIPA buffer [50 mM Tris-HCl (pH7.5), 500 mM NaCl, 1 mM EDTA, 1% NP40 (V/V), 0.1% SDS (W/V), 0.5% sodium deoxycholate (W/V), 1 mM DTT, 1X protease inhibitor cocktail]. 200 µL of streptavidin coated magnetic beads per sample (Catalog number: 65001, Dynabeads MyOne Streptavidin C1, Invitrogen, Vilnius, LT) were equilibrated by washing 5 times with RIPA buffer at 4°C. Equilibrated beads were incubated with the solubilized membrane fraction overnight at 4°C. Post incubation, the beads were harvested and sequentially washed with 1 mL buffer I (2% SDS in water) at room temperature, once with buffer II [50 mM HEPES (pH7.5), 500 mM NaCl, 1 mM EDTA, 0.1% deoxycholic acid (W/V), 1% Triton X-100], and once with buffer III [10 mM Tris-HCl, pH7.4, 250 mM LiCl, 1 mM EDTA, 0.1% deoxycholic acid (W/V), 1% NP40 (V/V)] at 4°C. Complete removal of the detergent was ensured by further washing the beads twice in 50 mM Tris-HCl, pH7.5 and six more times in 50 mM ammonium bicarbonate, pH 8.0. The beads resuspended in 1 mL 50 mM ammonium bicarbonate and flash-frozen in liquid nitrogen and submitted for LCMS analysis (Zhang *et al*., 2019; Sun *et al*., 2024). Proteins attached to beads were suspended in 20 µl of buffer containing 6 M urea, 2 M thiourea, 100 mM ammonium bicarbonate, and 5 mM DTT. Proteins were reduced at room temperature for 1 hr, then alkylated with 14 mM IAA for 30 minutes at room temperature in the dark and were digested with 0.4 µg of Trypsin overnight. The digestion was stopped adding TFA to a final concentration of 0.5%. One third of the digested peptides were used for protein quantification. EvoSep One liquid chromatography system was used for proteome analysis. The samples were processed with a 44-minute gradient for peptide elution (or Evosep One 30 SPD program). A 15 cm × 150 μm ID column with 1.5 μm C18 beads (Bruker PepSep) and a 20 µm ID zero dead volume electrospray emitter (Bruker Daltonik, GmbH, Germany) were used. Mobile phases A and B consisted of 0.1% formic acid in water and 0.1% formic acid in ACN, respectively. The Evosep One was coupled online to a modified trapped ion mobility spectrometry quadrupole time-of-flight mass spectrometer (timsTOF Pro 2, Bruker Daltonik, GmbH, Germany) via a nanoelectrospray ion source (Captive spray, Bruker Daltonik, GmbH, Germany). To perform data-independent acquisition (DIA-PASEF), the timsTOF Pro2 operated in a DIA-PASEF mode. The method employed 16 PASEF scans, each containing 4 overlapping ion mobility (IM) and m/z windows, resulting in 64 total precursor windows covering an m/z range of 400-1200 with narrow 25 m/z isolation windows and an ion mobility range of 0.6-1.6 Vs·cm⁻². The duty cycle was maintained at 100%, with a total cycle time of 1.8 seconds, consisting of one 100 ms MS1 survey scan followed by sixteen 100 ms DIA-PASEF scans. DIA-PASEF raw data was analyzed using the directDIA workflow in Spectronaut (version 20) with default settings. Trypsin was specified as the digestion enzyme, allowing up to two missed cleavages. Carbamidomethylation of cysteine was set as a fixed modification, while oxidation of methionine, N-terminal protein acetylation, and phosphorylation on serine, threonine, and tyrosine (STY) were included as variable modifications. Searches were performed against the *Arabidopsis thaliana* protein database. A false discovery rate (FDR) threshold of 1% was applied at the PSM, peptide, and protein levels for identification.

### Recombinant protein purification and *in vitro* pull-down

Full length PhyB coding sequence in frame with N-terminal 3x-HA, RBOHD N-terminus (aa1-376) and RBOHD C-terminus (aa756-922) in frame with N-terminal MBP were cloned in pDest-566 vector. Primers used to generate these constructs are listed in Table S5. To facilitate proper folding and solubilization of recombinant protein, individual constructs were co-transformed into BL21 *E. coli* competent cells with pG-KJE8 plasmid (Catalog number: 3340, Takara, San Jose, CA, USA). Cells were inoculated in 50 mL LB media in 250 mL conical flask and incubated on a shaker incubator at 37°C under 220 RPM until the OD at 600 nm reaches 0.5. Post incubation, flasks with the liquid culture were kept at 4°C for 20 minutes. Protein expression was induced by adding 0.5 mM isopropyl β-D-1-thiogalactopyranoside (IPTG), 2.5 mg/mL L-arabinose and 8 ng/mL tetracycline to the cultures and incubated further at 16°C under 220 RPM for 16 hours. Post incubation cells were harvested using centrifuge at 6,000 RPM for 10 minutes 4°C. Cell pellets were resuspended in 5 mL of lysis buffer [20 mM Tris-HCL at pH7.5, 200 mM NaCl, 1 mM EDTA, 0.1% triton, and 1X protease inhibitor cocktail (Lee *et al*., 2020)] and sonicated for 5 × 15 s with a 30 s interval between each sonication. Lysates were centrifuged at 14,000 RPM for 20 minutes 4°C. *E. coli* lysate with the bait protein MBP-RBOHD N-terminus (aa1-376) or MBP-RBOHD C-terminus (aa756-922) were incubated 2 hours at 4°C with the lysate expressing the HA-PhyB prey protein. *E. coli* lysate with the MBP protein served as control. The lysate mixture was then incubated overnight with 30 μL of equilibrated amylose resin (Catalog number: E8021S, New England BioLabs, Ipswich, MA, USA) at 4°C. Post incubation the beads were harvested and resuspended in 2X SDS loading buffer and ran on a 10% separating SDS gel. Western blot analysis was performed using anti-MBP (Catalog number; E8032S, New England BioLabs, Ipswich, MA, USA) or anti-PhyB antibody (Catalog number: AS214566, Vännäs, Sweden).

### Protein complex modeling and interaction hotspot annotation

Five Arabidopsis proteins FERONIA, PHYB, CRK10, RBOHD, and PIP2;6 were selected to evaluate their potential to form a complex. FASTA sequences and AlphaFold-predicted structures were retrieved from UniProt (UniProt Consortium, 2015) and the AlphaFold Database (Tunyasuvunakool *et al*., 2021). Domain boundaries were annotated using InterProScan (Jones *et al*., 2014) while functional motifs were identified using custom expression scans. Experimentally validated phosphorylation and post-translational modification (PTM) sites were compiled and integrated. Solvent accessibility (ASA/RSA) was calculated using FreeSASA (Mitternacht, 2016), and all annotations were merged into a unified hotspot table. Interface residues were prioritized based on their presence within annotated domains/motifs, overlap with PTMs, and solvent exposure. Pairwise protein docking was performed using the HDOCK (Yan *et al*., 2020) server to evaluate potential interaction surfaces. Docked complexes were visualized in PyMOL (Delano, 2002), and hotspot residues were highlighted with custom scripts to map annotated features onto structural models. For multimeric modeling, all five proteins were submitted to AlphaFold3 webserver (Krokidis *et al*., 2025), and the resulting complex was parsed and interpreted as five individual chains representing each protein. Subdomains of interest were extracted for focused visualization.

### Statistical analysis

Two-tailed Student’s t-test was used to determine the significance of variance (*P < 0.05) when comparing two treatments or genotypes. Analysis of variance (one way ANOVA/FISHER LSD) was used to determine the significance of variance (*P < 0.05) when comparing multiple genotypes and/or treatments with each other.

## AUTHOR CONTRIBUTIONS

D.M., Y.F., M.A.P.V., R.J.M.J., M.S., R.S., J.M., and R.E performed experiments and analyzed the data. D.M., E.L., Y.F., S.I.Z., and R.M. designed experiments. E.L., C.Y.Y., K.O., C.Z., and J-K.Z. provided essential research tools. C.X. and H.A. conducted *in-silico* modeling. D.M. M.A.P.V., and R.M. wrote the original draft of the manuscript. All authors approved the text and provided feedback. R.M. and E.L. provided financial support.

## ACKNOWLEDGMENTS

This work was supported by funding from the National Science Foundation (IOS-2414183; IOS-2110017, IOS-2343815) and the Interdisciplinary Plant Group, and University of Missouri, Columbia.

## DECLARATION OF AI USE

AI-assisted technologies were not used in creating this article.

## DATA AVAILABILITY

The data that supports the findings of this study are available in the text, figure, and supplemental material of this article. Proteomics data was deposited in PRIDE (https://www.ebi.ac.uk/pride/), under the following accession number: PXD070568.

## LIST OF SUPPLEMENTARY MATERIAL

**Figure S1.** Complementation of the *phyB* mutant with the phyB-miniTurbo protein.

**Figure S2.** *In vitro* protein-protein interaction studies between RBOHD (N-and C-terminals) and (PHYTOCHROME B).

**Figure S3.** Whole-plant reactive oxygen species (ROS) imaging analyses at time 0 of the PHYTOCHROME B mutant (*phyB*) complemented with wild type phyB or mutants of phyB.

**Table S1.** List of proteins that interact with the HA-RBOHD protein.

**Table S2.** List of proteins found in proximity to the phyB-miniTurbo protein.

**Table S3.** Summary of the Top 10 *In Silico* models for phyB interactions.

**Table S4.** Summary of the Top 10 *In Silico* models for RBOHD interactions.

**Tables S5.** Transcriptome-based tissue expression and predicted subcellular localization of all members of the putative complex identified in this study.

**Tables S6.** List of primers used in the study.

---

**Figure S1.**
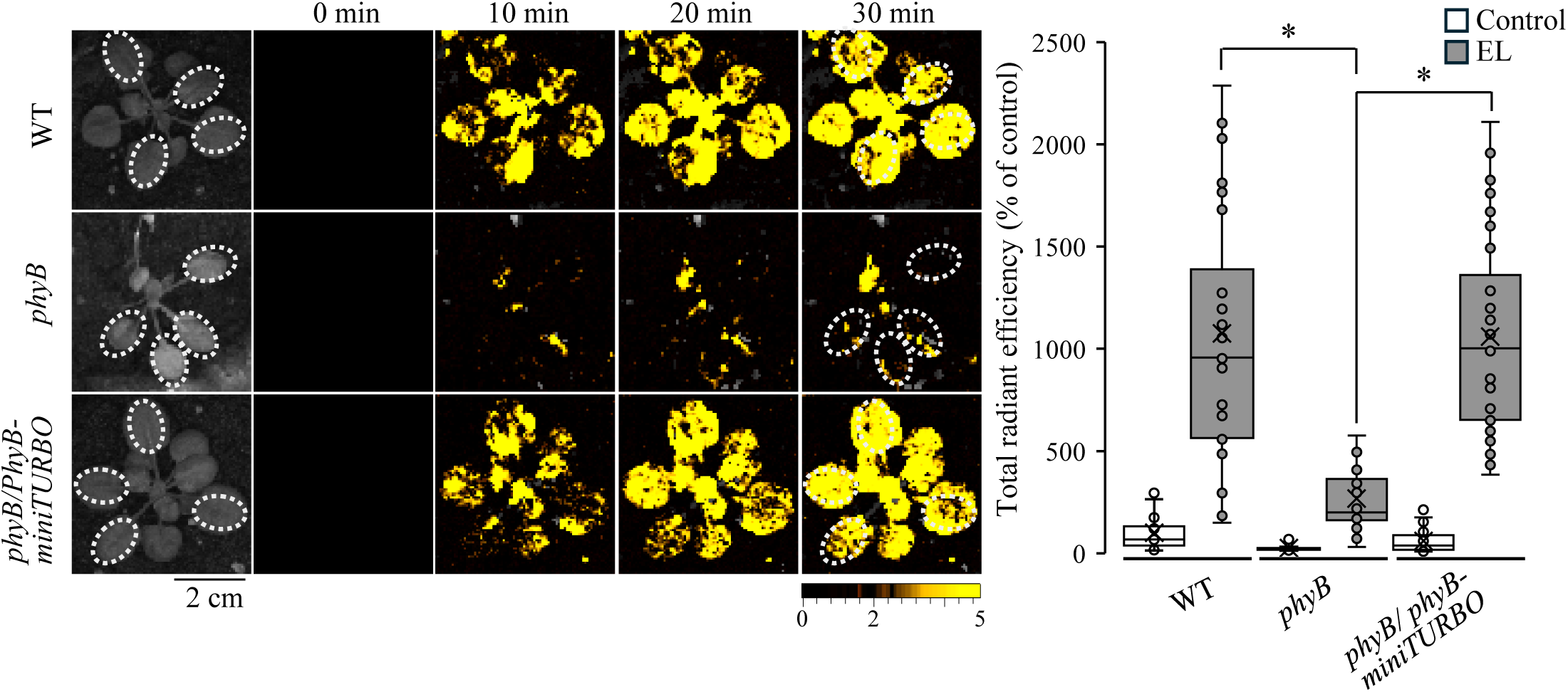
Complementation of the *phyb* mutant with the phyB-miniTurbo protein. Representative time-lapse whole-plant images (Left), and a bar graph showing quantification (Right) of reactive oxygen species (ROS) accumulation in wild type (WT), *phyb*, and *phyb* plants complemented with the phyB-miniTurbo protein driven by the ubiquitin 10 (*UBQ10*) promoter, in response to a whole-plant 10 min excess light (EL) stress. In support of Fig. 4a. Statistical significance was determined by using Student’s *t*-test: \**P* < 0.05 (N≥3). Abbreviations: cm, centimeter; GFP, Green Fluorescence Protein; min, minutes; *phyb*, PhytochromeB; WT, wild type.

**Figure S2.**
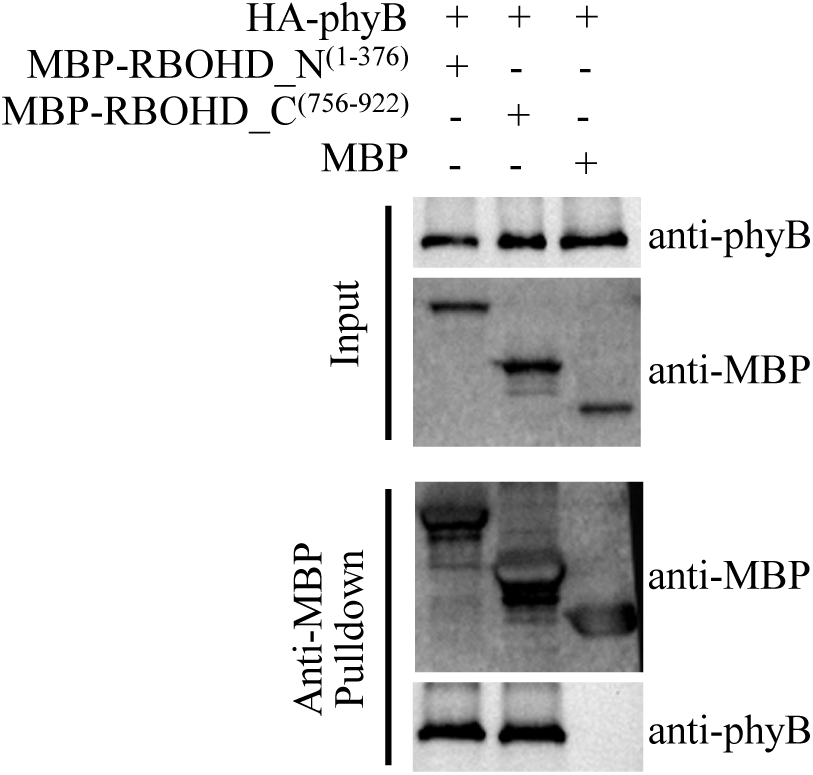
*In vitro* protein-protein interaction studies between RBOHD (N-and C-terminals) and (PHYTOCHROME B). *E. coli* expression vectors encoding HA-phyB, MBP-RBOHD-N (aa 1–376), MBP-RBOHD-C (aa 756–922), and MBP alone were generated, and recombinant proteins were expressed. Lysates containing HA-phyB were incubated with either MBP-RBOHD-N, MBP-RBOHD-C, or MBP, followed by pull-down using the MBP tag. HA-PhyB was detected in the pull-down fractions of both MBP-RBOHD-N and MBP-RBOHD-C, but not in the MBP control, providing additional evidence of phyB’s interaction with both the N-and C-terminal regions of RBOHD. In support of Figs. 4 and 5. Abbreviations: HA, hemagglutinin; MBP, Maltose Binding Protein; phyB, Phytochrome B; RbohD_C, Respiratory burst oxygen homologue D C-terminal; RbohD_N, Respiratory burst oxygen homologue D N-terminal.

**Figure S3.**
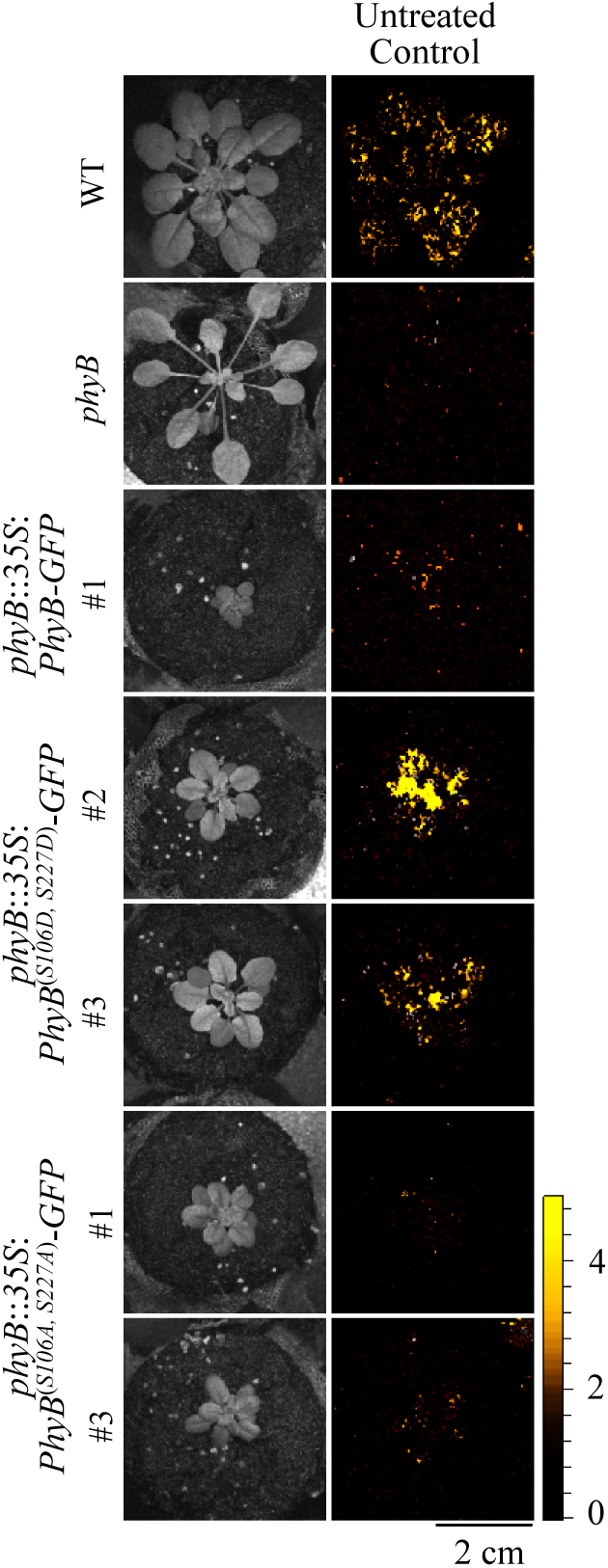
Whole-plant reactive oxygen species (ROS) imaging analyses at time 0 of the PHYTOCHROME B mutant (*phyB*) complemented with wild type phyB or mutants of phyB. Representative time-lapse whole-plant ROS images showing the ROS signal at time 0 for wild type (WT), the *phyb* mutant, the *phyb* mutant complemented with the phyB protein expressed under the control of the *CaMV35* promoter (*phyb*:3*5S*:*PhyB-GFP*), the *phyb* mutant complemented with a phyB protein that mimics constitutive phosphorylation of phyB [*phyb*:*PhyB^(S106D/S227D)^-GFP*], or the *phyb* mutant complemented with a phyB protein that cannot be phosphorylated by FERONIA [*phyb*:*PhyB^(S106A/S227A)^-GFP*]. In support of Fig. 6. Abbreviations: cm, centimeter; GFP, Green Fluorescence Protein; min, minutes; phyB, Phytochrome B; WT, wild type.

